# Alpha 6 integrins promote cytokinesis by regulating the expression of RSK2 and MKLP1

**DOI:** 10.1101/830562

**Authors:** Neetu Singh, Hao Xu, Renee Thiemann, Kara A. DeSantis, Melinda Larsen, Susan E. LaFlamme

## Abstract

The integrin-mediated interaction of cells with components of the extracellular matrix (ECM) regulates many cellular processes including cell division. Cytokinesis is the last step of cell division and is critical for normal development and tissue homeostasis as it ensures the proper segregation of genetic and cytoplasmic material between daughter cells. Cytokinesis failure leads to defects in development and tissue differentiation, as well as tumorigenesis. Abscission of intercellular bridge that connects presumptive daughter cells is the last step of cell division. The mitotic kinesin-like protein 1 (MKLP1) plays a central role in positioning the abscission machinery. Here, we show that α6 integrins promote successful cytokinesis in salivary gland epithelial cells by regulating the expression of MKLP1. RNAi-mediated depletion of α6 integrins inhibits cytokinesis and the expression of MKLP1 and p90 ribosomal-S6-kinase 2 (RSK2). Depletion of RSK2 results in similar defects in cytokinesis and also inhibits the expression of MKLP1, suggesting that the expression of RSK2 is required downstream of integrins to promote MKLP1 expression and successful cytokinesis. RNAi-mediated depletion of RSK2 in embryonic salivary glands in organ culture also results in the inhibition of cytokinesis and MKLP1 expression, indicating the physiological significance of this pathway.

## INTRODUCTION

Integrins are α/β transmembrane receptors that mediate cell adhesion to the ECM (1). Individual cell types express different sets of integrin α/β heterodimers to bind to distinct components of the ECM. Laminins are the major ECM components expressed by epithelial cells. Alpha6 integrins, the α6β1 and α6β4 heterodimers, are one class of integrins expressed by epithelial cells that mediate adhesion to laminin (1). Integrin-dependent interaction of cells with components of the ECM, including laminin, activates signaling pathways to control many aspects of cell behavior, such as migration, survival, gene expression, differentiation, and proliferation, including cytokinesis (2–4).

Cytokinesis is the last step of cell division and is critical for normal development and tissue homeostasis as it ensures the proper segregation of genetic and cytoplasmic material between daughter cells (5). Cytokinesis begins with the assembly and constriction of the contractile ring that drives the ingression of the cleavage furrow, and concludes with abscission of the intercellular bridge (as referred to as the midbody) that connects presumptive daughter cells (5). Defective or failed cytokinesis promotes tumorigenesis (6–8), whereas the deletion of genes required for cell division blocks embryonic development (9,10). Integrins impact these processes, as the loss of integrin expression or activity can lead to defective cytokinesis, resulting in defective tissue differentiation (11,12) or aneuploidy and tumor formation (13).

Much of what is known about the mechanisms that link integrins and cytokinesis have been identified using cell culture models. Our published studies identified a role for integrins in the regulation of abscission (14) and have defined a role for ERK and its downstream target the ribosomal S6 kinase (RSK) (14). RSKs are a family of serine/threonine kinases that phosphorylate many cytoplasmic and nuclear targets (15–17). We previously demonstrated that depletion of either RSK1 or RSK2 in mammary epithelial cells inhibited cytokinesis (14), but did not determine whether both RSK isoforms were activated downstream of integrins or whether RSK1 and RSK2 regulated cytokinesis by similar or distinct mechanisms. Our current study addresses these gaps in knowledge and further explores the important role of RSK2 in the successful completion of cytokinesis.

Abscission of the intercellular bridge/midbody is catalyzed by the endosomal-sorting complex required for transport (ESCRT), together with additional proteins (18). The ESCRT complex is recruited to the midbody by stepwise assembly of proteins beginning with the mitotic kinesin-like protein1, MKLP1, then the centrosomal 55 kD protein (CEP55), and Alg2-interacting protein X (ALIX) followed by the ESCRT complex (5). Here, we tested whether α6 integrins promote cytokinesis by regulating the expression of any of these midbody components and whether RSK1 and/or RSK2 were required for this regulation.

To test this, we first used SIMS cells (19,20), a murine salivary gland epithelial cell line that allowed us to easily examine mechanisms that regulate cytokinesis downstream of α6 integrins employing a cell culture model. We then confirmed the physiological significance of these mechanisms using the murine salivary gland in organ culture. We demonstrate that integrins regulate the expression of MKLP1 and that RSK2 functions downstream of integrins in the regulation of MKLP1 expression and successful cytokinesis. We show that RSK1 and RSK2 regulate cytokinesis by distinct mechanisms and that only RSK2 functions downstream of α6 integrins. Lastly, we show that RSK2 also regulates cytokinesis and the expression of MKLP1 in *ex vivo* organ culture, indicating that this pathway is important not only in cell culture, but also in the more complex physiological context of the embryonic murine salivary gland.

## RESULTS

### Alpha 6 integrins regulate MKLP1 expression

To determine whether α6 integrins regulate the expression of proteins required for the assembly of the abscission machinery, we inhibited the expression of the integrin α6 subunit using RNAi technology targeting different regions of the α6 mRNA. We determined the effect of α6 depletion on the expression of MKLP1, CEP55, and ALIX, which are proteins involved in the recruitment of the ESCRT complex. As an initial approach, we generated a lentiviral vector for the doxycycline inducible expression of an α6 targeting shRNA. SIMS cells infected with the lentivirus were selected and treated with or without doxycycline. The induction of the α6-targeting shRNA significantly inhibited α6 expression compared to control (Fig. 1A). Depletion of α6 integrins resulted in the inhibition of cytokinesis. We observed a significant increase in the number of cells connected by midbodies, indicative of a delay in abscission, and the accumulation of binucleated cells resulting from failed cytokinesis (Figs. 1B-C). The shRNA-dependent depletion of α6 integrins also results in a significant decrease in the expression of the MKLP1 protein when assayed by western blot and quantified from three independent experiments (Fig. 1D-E). Additionally, qPCR analysis indicated MKLP1 transcripts levels were also significantly decreased in α6-depleted cells (Fig.1F). In contrast, the expression of CEP55 and ALIX were not affected (Fig. 1D-F). Analogous results were obtained using an siRNA that targeted a different region of the α6 mRNA (Fig. 1. G-L). Thus these results indicate that α6 integrins regulate RNA and protein levels of MKLP1, a key cytokinesis regulator.

**Fig. 1.**
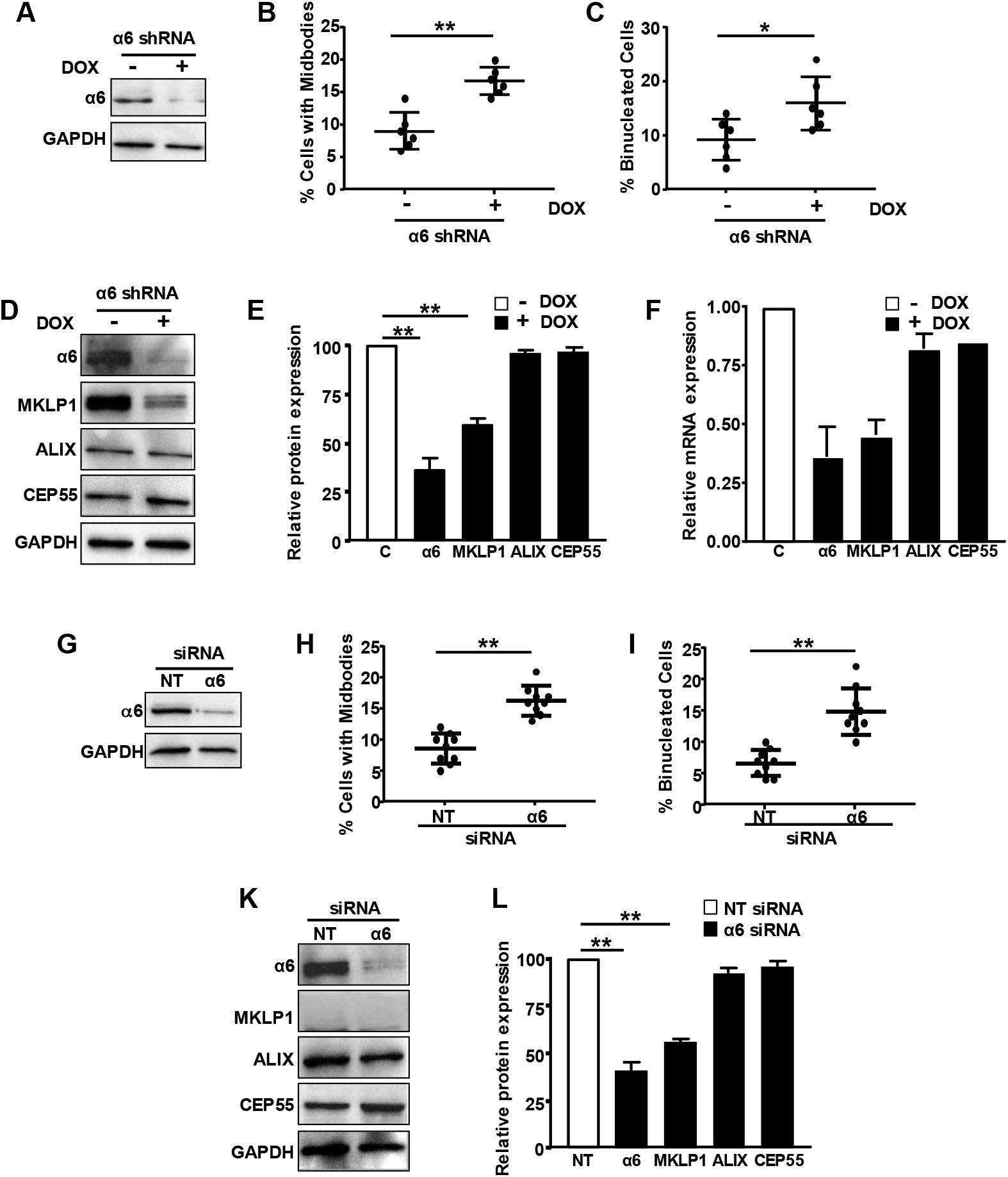
Depletion of α6 integrins inhibits cytokinesis and the expression of MKLP1. **(A)** SIMS cells harboring an inducible shRNA targeting the integrin α6 subunit were treated with doxycycline to induce the expression of the α6 shRNA or left untreated. The efficiency of α6 knockdown was assayed by western blot. Shown is a representative blot probed with antibodies to the α6 integrin subunit and reprobed with antibodies to GAPDH as a loading control. **(B)** Cells expressing the shRNA or control cells were stained for α-tubulin and DNA to quantify cells connected by midbodies and **(C)** binucleated cells. Three hundred individual cells were analyzed from each of three independent experiments performed in triplicate. Plotted is the mean percentage of cells with mibodies or that are binucleated ± s.d. (n = 9). **(D)** Effect of α6 depletion by shRNA on the expression of midbody proteins required for cytokinesis was assayed by western blot. Shown is a representative blot. **(E)** Quantitation from three independent experiments ± s.d. **(F)** The effect of α6 depletion by shRNA on mRNA expression was assayed by qPCR. Data is shown from two independent experiments ± variance. **(G-L)** SIMS cells were transfected with either NT or α6 targeting siRNA. The effect of α6 depletion on cytokinesis **(G-I)** and midbody components was assayed as above for α6 targeting shRNA **(K-L). (K)** Shown is a representative blot and **(L)**quantitation of protein expression from three independent experiments. **p<0.05, *p<0.001.

### RSK2 functions downstream of α6 integrins

Although our previous studies demonstrated that both RSK1 and RSK2 regulated cytokinesis in MCF10A cells (14), these studies did not determine whether both isoforms functioned downstream of integrins or whether they regulated cytokinesis by similar mechanisms. To begin, we confirmed that both RSK1 and RSK2 regulated cytokinesis in SIMS cells. We inhibited their expression using RSK1 and RSK2 targeting siRNAs (Fig. 2A-B). As expected, depleting SIMS cells of RSK1 or RSK2 delayed abscission as indicated by an increase in cells connected by midbodies and led to the formation of binucleated cells (Fig. 2A-B).

**Fig. 2.**
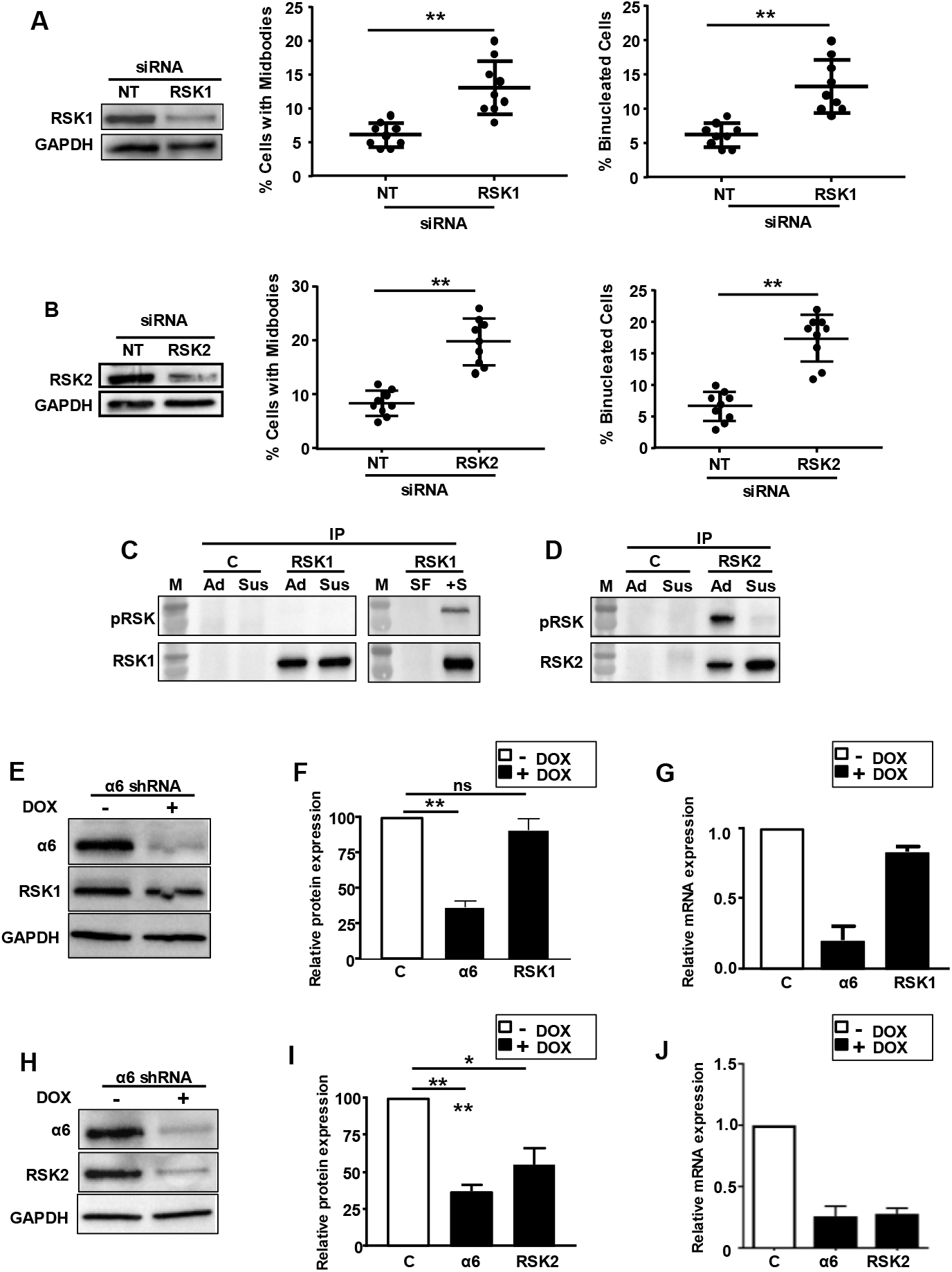
Depletion of RSK1 or RSK2 inhibits cytokinesis, but only RSK2 functions downstream of integrins. **(A-B)** SIMS cells were transfected with non-targeting, RSK1 targeting or RSK2 targeting siRNA. The efficiency of knockdown of RSK1 **(A)** and RSK2 **(B)** was analyzed by western blotting (left panels). The effect of depletion on cytokinesis was determined by monitoring the persistence of midbodies (middle panels) or binucleated cells (right panels). Plotted is the mean percentage of cells in ± s.d. Three hundred individual cells were analyzed from each of three independent experiments performed in triplicate (n = 9). **(C-D)** Adhesion-dependent activation of RSK1 and RSK2 was determined by incubating cells either in suspension (Sus) or re-adhered (Ad) to a laminin matrix. **(C)** RSK1 and **(D)** RSK2 were immunoprecipitated with isoform specific antibodies and analyzed for RSK activation by western blot using an activation-specific antibody (pRSK) and then reprobed for RSK1 **(C)** or RSK2 **(D)**. Although RSK1 was not activated upon adhesion it was activated by the addition of serum (S) compared to serum free medium (SF) **(C). (E-F)** The effect of depletion of α6 on the expression of RSK1 **(E)** and RSK2 **(H)** was analyzed by western blotting. Protein expression was quantified from three independent experiments **(F&I)**. The effect of α6 depletion on RSK1 **(G)** and RSK2 **(J)** RNA levels was analyzed by qPCR. Plotted is mean ± variance from two independent experiments. *p<0.05, **p<0.001.

Since both RSK1 and RSK2 inhibited cytokinesis in SIMS cells, we were interested to determine whether α6 integrins differentially regulated the activation or expression of RSK1 and RSK2. We first tested whether cell adhesion to laminin, a known substrate for α6 integrins resulted in the activation of RSK1 and/or RSK2. SIMS cells were either incubated in suspension or re-adhered to a laminin matrix for 2.5 h. RSK1 and RSK2 were then separately immunoprecipitated using isoform specific antibodies and RSK activation was evaluated by western blot using a phosphorylation-specific antibody that recognizes active RSK (15). RSK1 was not activated by adhesion; however, RSK1 was activated in response to the addition of serum (Fig. 2C). In contrast, RSK2 was activated when cells were adhered to laminin when compared with cells that were deprived of adhesion (Fig. 2D).

To determine whether α6 integrins differentially regulate the expression of RSK isoforms, SIMS cells were depleted of α6 using the inducible expression of α6 targeting shRNA as described above. The effect on RSK expression was analyzed by western blot and qPCR. The results indicate that depleting cells of α6 integrins did not affect the levels of RSK1 protein, as analyzed by western blot and quantified from three independent experiments (Fig. 2E-F). This was also true of RSK1 transcript levels (Fig. 2G). In contrast, depletion of α6 integrins led to the inhibition of RSK2 levels both at the protein (Fig. 2H-I) and mRNA level (Fig. 2J). Taken together our data suggest that RSK2 functions downstream of α6 integrins to promote successful cytokinesis.

### RSK2 regulates the expression of MKLP1

As shown above, depleting cells of α6 integrins decreases the levels of RSK2, as well as MKLP1. This suggests that RSK2 may regulate cytokinesis by regulating the expression of MKLP1. To test this, SIMS cells were depleted of RSK2 using RNAi and the effect on the expression of MKLP1, CEP55, and ALIX was analyzed by both western blot and qPCR. Depleting cells of RSK2 inhibited the expression of MKLP1 protein, as analyzed by western blot and quantified from three independent experiments (Fig. 3A-B). As we observed with the depletion of α6 integrins, inhibiting the expression of RSK2 had no effect on the expression of CEP55 and ALIX (Fig. 3A-B). Additionally, qPCR analysis indicated that RSK2 depletion also inhibited the expression of MKLP1 transcripts (Fig. 3C). Similar results were obtained when RSK2 expression was inhibited using an inducible shRNA targeting RSK2 (Fig. 2D-E). In contrast, depleting cells of RSK1 did not affect the levels of MKLP1 (Fig. 1F). Taken together our results suggest that α6 integrins regulate cytokinesis by regulating the RSK2-dependent expression of MKLP1 and indicate that RSK1 and RSK2 regulate cytokinesis by independent mechanisms.

**Fig. 3.**
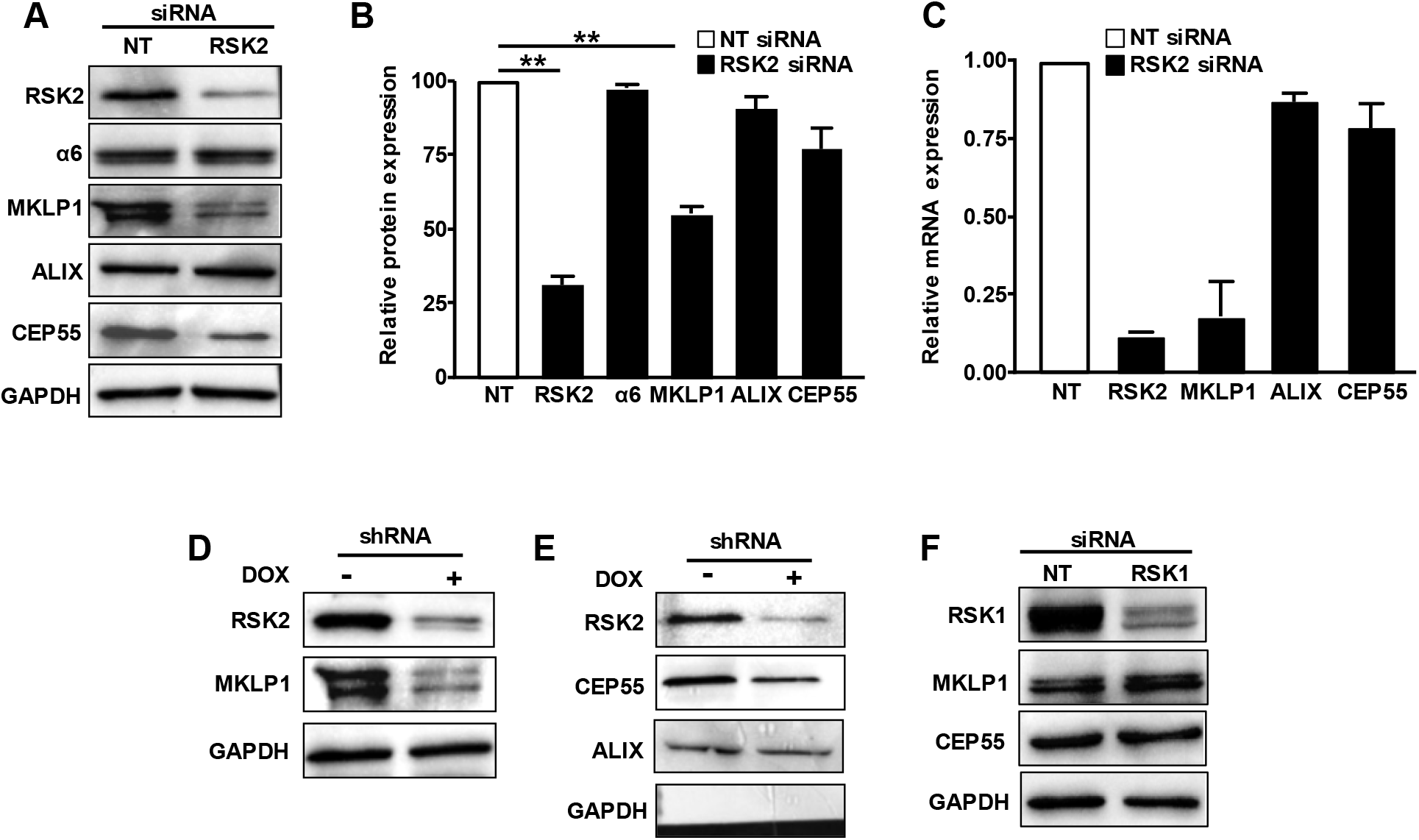
RSK2 inhibits the expression of MKLP1 protein and mRNA transcripts. SIMS cells were transfected with RSK2 or RSK1 siRNAs. **(A-B)** The effects of RSK2 depletion on α6 and MKLP1 and other cytokinesis proteins were assayed by western blot. **(A)** Shown is a representative blot and **(B)** quantitation from three independent experiments ± s.d. **(C)** The effect of RSK2 depletion on mRNA expression was assay by qPCR. Plotted is the mean ± variance from two independent experiments. **(D-E)** Inhibiting the expression of RSK2 by a doxycyline inducible shRNA results in similar effects and inhibited the expression of MKLP1 protein and transcripts **(F)** The siRNA-dependent depletion of RSK1 did not affect the expression of MKLP1. **p<0.001

### Depletion of MKLP1 results in similar cytokinesis defects as the depletion of α6 integrins or RSK2

MKLP1 is a critical regulator of cytokinesis and previous studies have demonstrated that RNAi-dependent depletion of MKLP1 results in cytokinesis failure (21). Since we propose that α6 and RSK2 regulate the expression of MKLP1 to promote cytokinesis, we tested whether the depletion of MKLP1 resulted in a similar level of cytokinesis failure in SIMS cells. To test this, we transfected SIMS cells with siRNA targeting MKLP1. Depletion of MKLP1 resulted in a similar level of cytokinesis failure as depletion of either α6 or RSK2 (Fig 4. A-C). These results support a mechanism in which α6 integrins positively regulate the expression of RSK2, which in turn positively regulates the expression of MKLP1 to promote cytokinesis.

**Fig. 4.**
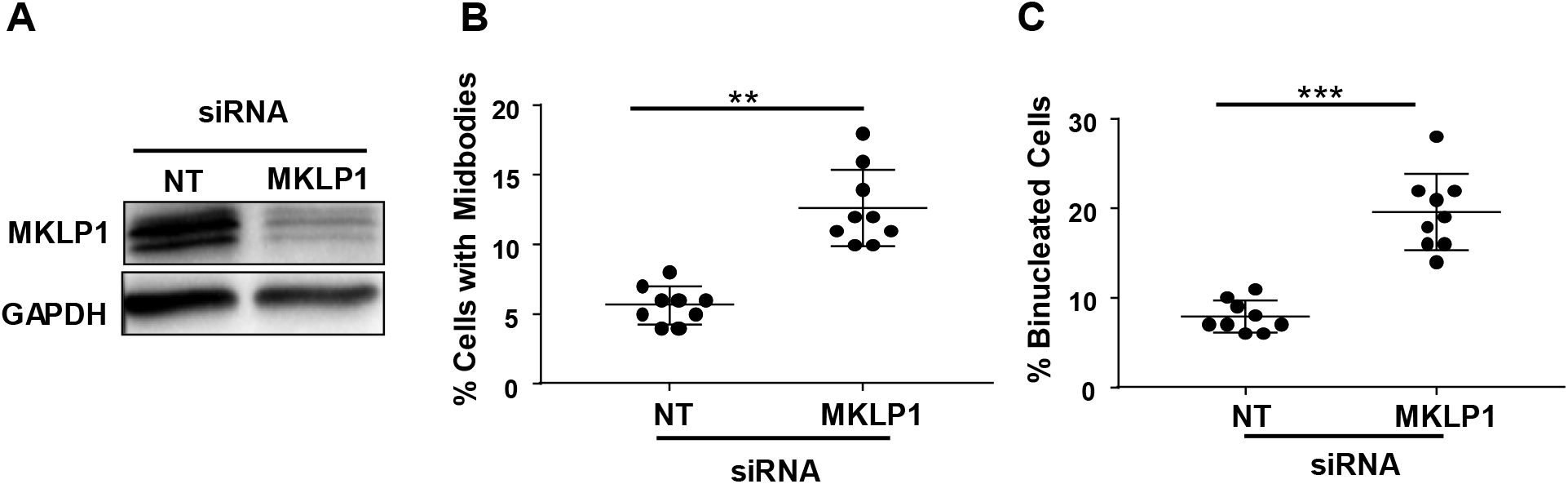
Depletion of MKLP1 inhibits cytokinesis in SIMS cells. **(A-C)** SIMS cells were transfected with MKLP1 siRNA. **(A)** The efficiency of depletion was determined by western blot. The effect of MKLP1 depletion of cytokinesis on the persistence of midbodies and the formation of binucleated cells was analyzed. Plotted is the mean percentage of cells with midbodies **(B)** or that are binucleated **(C)** ± s.d. Three hundred cells were analyzed from each of three independent experiments performed in triplicate. ** p<0.001, ***p<0.0001.

### RSK2 regulates the expression of MKLP1 in the embryonic salivary gland

Much of what is known about the mechanisms that regulate cytokinesis in mammalian cells has been acquired from experiments in culture using cell lines that are easily transfected and imaged (22). However, since integrin signaling can be strongly influenced by substrate compliance (23), and our previous studies demonstrated that cytokinesis is regulated by matrix compliance in a cell type-dependent manner (24), it is important to demonstrate that pathways that regulate cytokinesis in cell culture similarly regulate cytokinesis in a more physiological complex context. Since we previously demonstrated that α6 integrins promote successful cytokinesis in embryonic salivary glands in *ex vivo* organ culture (14), we again used this model to test whether RSK2 regulates cytokinesis and MKLP1 expression in this context. To this end, submandibular salivary glands were isolated from mouse embryos at embryonic day 12.5-13 (E12.5-E13) and treated for 3 days with either nontargeting or RSK2-targeting siRNA (Fig. 5A). The effects on RSK2 expression and cytokinesis were analyzed from three independent experiments. The efficiency of RSK2 knockdown was analyzed by western blot and quantified from the three experiments (Fig. 5B), which demonstrated that RSK2 depletion significantly inhibited cytokinesis. The results indicate that ~25% of cells from glands treated with RSK2 siRNA were binucleated compared with ~8% of cells from glands treated with the non-targeting siRNA (Fig. 5C). Importantly, RSK2 depletion also inhibited the expression of MKLP1 in this context, as analyzed by western blot from two independent experiments (Fig. 5D-E). Thus, RSK2 regulates cytokinesis by similar mechanisms both in salivary gland epithelial cells in culture and in embryonic salivary glands in organ culture.

**Fig. 5.**
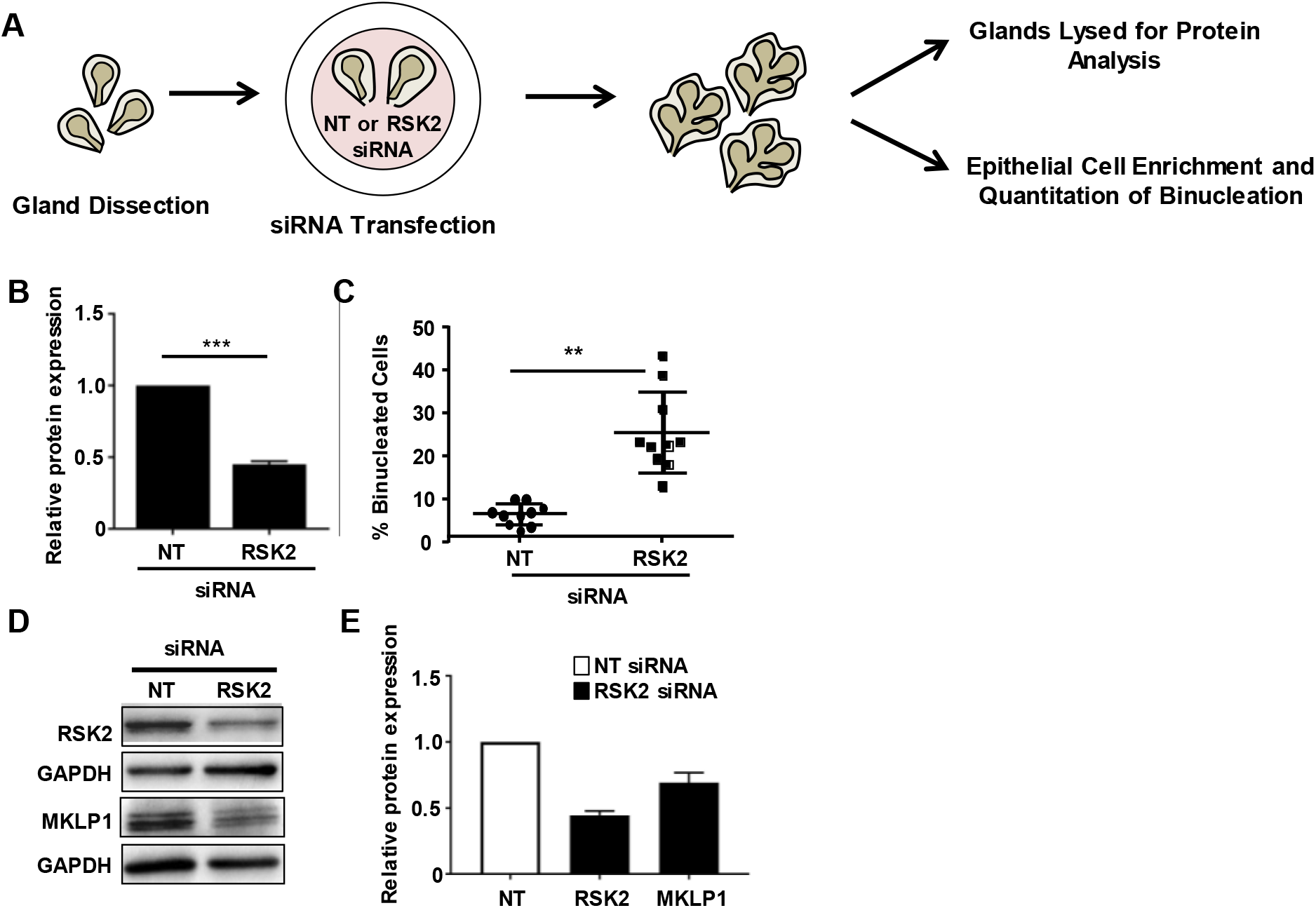
Depletion of RSK2 inhibits cytokinesis and MKLP1 expression during salivary gland morphogenesis in *ex vivo* organ culture. **(A)** Shown is a schematic of the experimental design. **(B-C)** Submandibular salivary glands were treated with RSK2-targeting or non-targeting siRNA starting at E12.5. **(B)** The efficiency of RSK2 knockdown was assayed by western blot and **(C)** quantified from three independent experiments to determine the effects on cytokinesis. Cytokinesis was assayed in 15 glands from each treatment, and approximately, 100 cells were analyzed per sample from each of three independent experiments. Plotted is the mean percentage of binucleated cell ± s.d. **(D)** Representative western blots of MKLP1 expression in glands treated with NT- or RSK2-targeting siRNA. **(E)** Quantitation of data from 2-independent experiments plotted as the mean ± variance. **p<0.001, ***p<0.0001.

## DISCUSSION

In this study, we demonstrated that α6 integrins promote successful cytokinesis in cultured salivary gland epithelial cells, at least in part, by the α6-dependent expression of MKLP1, a key regulator of cytokinesis. Our data also show that α6 integrins promote the expression of RSK2 and that RSK2 positively regulates the expression of MKLP1. We propose that α6 integrins promote a gene expression cascade in which α6 integrins regulate cytokinesis by promoting the expression of RSK2, which in turn promotes the expression of MKLP1 (Fig. 6). Additionally, our results indicate that although the depletion of either RSK1 or RSK2 inhibits cytokinesis in salivary gland epithelial cells, only RSK2 is regulated by α6 integrins and that RSK1 and RSK2 regulate cytokinesis by distinct mechanisms.

**Fig. 6.**
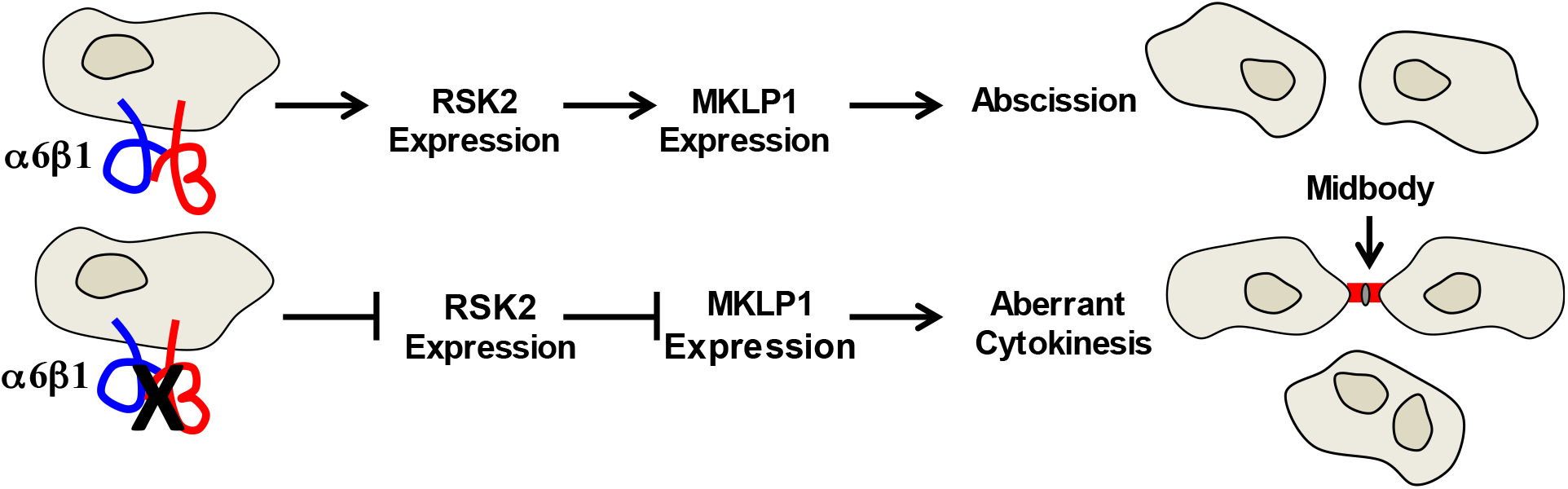
Model for the regulation of cytokinesis by α6 integrins.

It is well established that the compliance of the adhesion substrate impacts integrin signaling and cytokinesis (23,24). Our studies with SIMS cells were performed on culture plates, whose stiffness is orders of magnitude higher than observed in tissues (25). Therefore, it was critical to demonstrate a role for RSK2 in a more physiological context, allowing epithelial cells to interact with the ECM in a physiological compliant environment. We accomplished this by inhibiting the expression of RSK2 in embryonic salivary glands *ex vivo* organ culture. Depletion of RSK2 resulted in the inhibition of MKLP1 expression and impaired cytokinesis, confirming the significance of this pathway in a physiological context.

In addition to regulating the expression of MKLP1, our previous studies suggested that RSK activity is also required proximal to abscission, as the pharmacological inhibition of RSK after the formation of the midbody/intercellular bridge was sufficient to inhibit cytokinesis (14). We and others, have observed the localization of RSK2 in the midbody/intercellular bridge of dividing cells ((26) and N. Singh (unpublished data)), suggesting that RSK2-dependent phosphorylation of a component of the abscission machinery may promote the successful completion of cytokinesis. Thus, RSK2 may regulate cytokinesis by regulating both the expression and activity of components of the abscission machinery. RSK2 activity may also be required at late stages of cytokinesis to modulate integrin activity. Active RSK2 has been shown to down regulate integrin activity (27). Thus, the RSK2-dependent modulation of integrin activity may be required for abscission, as studies have shown that tension exerted on the intercellular bridge by presumptive daughter cells must be released for membrane fission to occur (28).

Our previous studies demonstrated that RSK2 does not regulate cytokinesis in human fibroblasts (14), indicating the regulation of cytokinesis by RSK2 exhibits cell type specificity. Interestingly, others have identified a role for FAK-SRC signaling in promoting cytokinesis of human fibroblasts (29). The pharmacological inhibition of FAK or SRC was shown to disrupt the appropriate timing of the recruitment of CEP55 to the midbody (29), which is known to result in abscission failure (30). The ability of Src signaling to promote abscission does not appear to be cell type specific, as blocking Src signaling inhibited abscission in Hela cells (31). Hence, integrins promote cytokinesis by cell-type specific and common signaling pathways.

Multiple integrin heterodimers can promote successful cytokinesis and do so in an integrin ligand-dependent manner (14,32–35). Integrin trafficking to the cleavage furrow is important as it supports the adhesion of the cleavage furrow to the underlying matrix (34). Some mechanisms regulating integrin trafficking rely on protein interactions with the integrin β1 cytoplasmic tail, and thus, impact cytokinesis in cells adhered to the multiple ligands for β1 integrins (34). However, other mechanisms are specific to individual integrin heterodimers. This is the case for the α5β1 heterodimer (35). The interaction of ZO-1 with the α5 subunit cytoplasmic domain is required for the trafficking of α5β1 and cytokinesis (35). Thus, integrin-regulated cytokinesis occurs by mechanisms shared by multiple integrin heterodimers and other specific to individual integrins.

In conclusion, we have identified α6 integrins and RSK2 as positive regulators of cytokinesis in epithelial cells, thus, protecting epithelial tissue homeostasis, as failed cytokinesis in epithelial cells has been reported to promote tumorigenesis (6–8).

### Experimental procedures Antibodies and Reagents

Mouse monoclonal antibody to α-tubulin (DM1α), goat polyclonal antibody to RSK2 (C-19), rabbit polyclonal antibody to CEP55 (M-300), mouse monoclonal antibody to the integrin α6 subunit (F6), and rabbit polyclonal antibodies to RSK1 (C21) were from Santa Cruz Biotechnology Inc. (Santa Cruz, CA). Rabbit monoclonal antibody to MKLP1 (EPR10879) was from Abcam (Cambridge, UK). Mouse monoclonal antibodies to ALIX (3A9) and GAPDH (clone GAIR) were from Thermo Fisher Scientific (Agawam, MA, USA). Rabbit polyclonal antibodies to phosphorylated serine 380 of RSK and DRAQ5 were from Cell Signaling Technology (Danvers, MA, USA). Alexa-Fluor-conjugated secondary antibodies and Hoechst 33342 were from Invitrogen. For immunofluorescence, the monoclonal antibody for α-tubulin and it corresponding secondary antibody were used at 1:800. For western blotting, antibodies were diluted to 1:500 except for antibodies to MKLP1 and pRSK, which were diluted to 1:1000.

### Cell Culture

Cell lines were cultured at 37°C in 5% CO_2_. The murine submandibular salivary gland epithelial cell line, SIMS cells (Laoide, 1996 and 1999), was cultured in DMEM/F12. The 804G bladder carcinoma cells (36) used to isolate laminin-332 (laminin-5) matrix and 293FT cells used in preparation of lentiviruses were cultured in DMEM. All culture media also contained 10% FBS, 100 units/ml penicillin, 100 μg/ml streptomycin and 2.92 ug/ml L-glutamine.

### SiRNA transfection in cell culture

SIMS cells were plated in 6 well tissue culture plates and treated with siRNA (80 nM per well) using Oligofectamine (Invitrogen/ThermoFisher) following the protocol provided by the manufacturer. After 3 days, the cells were trypsinized and a fraction of the cells was used for western blot analysis. The remainder were replated on coverslips, incubated overnight and then analyzed for cytokinesis failure as described below. SiRNAs, targeting and non-targeting, were purchased from Dharmacon/Thermo Scientific. The targeting sequences are the following: RSK1 siRNA: AGGGCAAGCUCUAUCUUAU; RSK2 siRNA: GCACGAAUAGGUAGUGGAA; α6 siRNA:GGACCAAAGACUCGAUGUU. A SMART pool was used in the case of MKLP1 (Dharmacon, Lafayette, CO).

### Generation of lentiviral vectors for shRNA expression

Sequences encoding shRNA for RSK1, RSK2 and α6 shRNA were cloned into the lentiviral vector, pLKO-Tet-On (Addgene plasmid # 21915) (37). The oligonucleotides encoding shRNAs were hybridized and then cloned into the AgeI and EcoR1 sites of pLKO-Tet-On lentiviral vector. Insertion of the correct sequences was confirmed by nucleotide sequence analysis. The oligonucleotides used in vector construction are as follows: <u>RSK2 oligonucleotides</u>: 5’-CCGGGCCGTGAAGATTATTGATAAACTCG AGTTTATCAATAATCTTCACGGCTTTTTG-3’ and 5’-CGGCACTTCTAATAACTATTTGACTCAAATAGTTATTAGAAGTGCCGAAAAACTTAA-3’. <u>α6 oligonucleotides</u>: 5’-CCGGCGTCTGATAAAGAGAGGCTTACTCGAGTAAGC CTCTCTTTATCAGACGTTTTTG-3’ and 5’-GCAGACTATTTCTCTCCGAATGAGCTCATT CGGAGAGAAATAGTCTGCAAAAACTTCAA-3’. <u>RSK1 oligonucleotides</u>: 5’-CCGGGCTCTAT CTTATTCTGGACTTCTCGAGAAGTCCAGAATAAGATAGAGCTTTTTG-3’ and 5’-CGAGA TAGAATAAGACCTGAAGAGCTCTTCAGGTCTTATTCTATCTCGAAAAACTTAA-3’.

Lentiviruses were produced by cotransfection of 293FT cells with the expression vector together with pCMV-dR8.2 and pCMV-VSV-G and then used to infect SIMS cells. Positively infected cells were selected with 2 *μ*g/ml puromycin (Gibco/Life technologies). Cells were removed from puromycin 48 h prior to shRNA induction with doxycycline (100 ng/ml).

### Western blotting

Western blotting was used to confirm siRNA and shRNA knockdown and to assay the effect of RSK2 depletion on various abscission components. Cells were lysed in mRIPA buffer containing phosphatase (Fisher Scientific) and protease inhibitor cocktails (Krackeler Scientific) and equal amounts of protein (30-40 μg) were separated by SDS-PAGE and transferred to nitrocellulose and then incubated with appropriate primary and secondary antibodies. Membranes were blocked in 2% BSA/PBST for 1 h and then incubated with primary antibody application overnight at 4°C. Membranes were washed in PBST prior to secondary antibody application. Membranes were exposed to SuperSignal West Pico PLUS Chemiluminescent substrate (Thermo Scientific, #34580) and SuperSignal West Femto Maximum Sensitivity substrate (Thermo Scientific, #34095) prior to imaging on a BioRad ChemiDoc™ MP Imaging System and analysis with Image Lab software. Protein signals were normalized to GAPDH.

### Immunoprecipitation

SIMS cells were isolated from logarithmically growing cultures and either replated on a laminin-rich matrix (38,39) or incubated in suspension for 2.5 h in CCM1 medium (Thermo Scientific). After incubation, the cells were washed with 1X PBS and solubilized for 15 min in 500 μl of lysis buffer (1% Nonidet P-40, 50 mM Tris-HCl (pH 7.4), 0.25% Deoxycholate (Na), 150 mM NaCl, 1mM EDTA, 1X protease inhibitor cocktail (Roche Applied Science), and 1X phosphatase inhibitor (PhosSTOP, Roche) on ice. Subsequent steps were performed at 4°C. Cell extracts were centrifuged for 5 min at 3000 rpm, and the supernatant was incubated for 4 h with corresponding control IgG and RSK antibody (4 *μ*g/ml), and then 45 *μ*l of protein A/G Plus Agarose beads (Pierce Biotechnology, Rockford, IL,) was added to each sample and samples were incubated for 45 min. Agarose beads/antibody complexes were recoverd by centrifugation, washed five times with wash buffer (1X PBS with 1% NP40). Proteins were solubilized in SDS-PAGE sample buffer and analyzed by western blotting.

### Quantitative PCR (qPCR)

RNA was isolated from SIMS cells and *ex vivo* cultures of murine submandibular salivary glands using Perfect RNA Cell Culture Kit-50 (5 Prime, Gaithersburg MD). cDNA was synthesized using iScript Reverse Transcription Supermix (BioRad) using 1 μg of DNase-treated RNA. Equal amounts of cDNA were used in qPCR reactions performed with iQSYBER Green Supermix (BioRad). Primer sequences are provided in Table I and were identified using the Harvard PrimerBank (40–42). Ct values were normalized to β-actin. Reactions were run in duplicate in 3 independent experiments.

**Table I.**
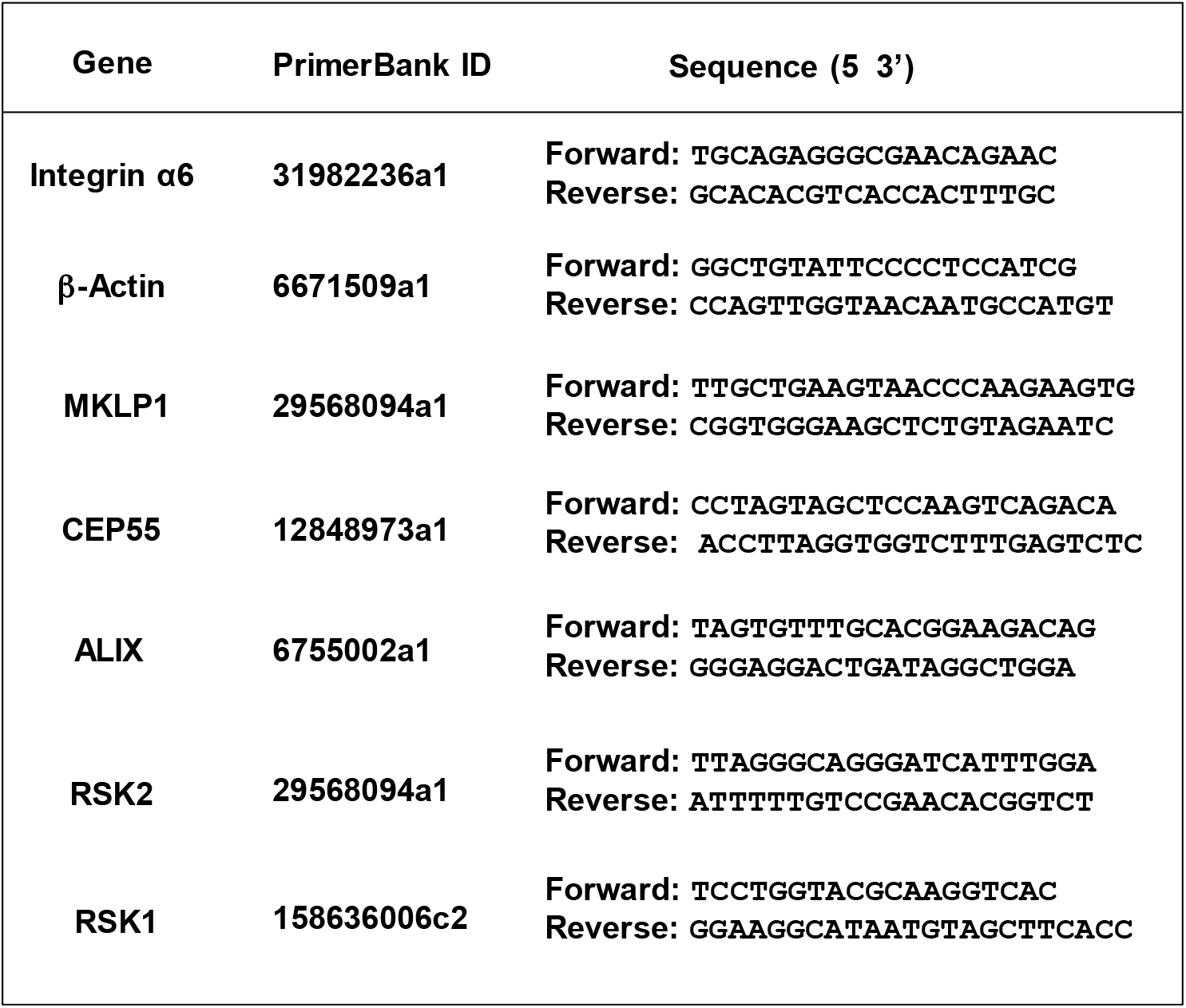
qPCR Primers

### Immunofluorescence

Cells were cultured either on glass coverslips and fixed in 4% paraformaldehyde for 20 min or icecold methanol for 5 min. Cells were then permeabilized with 0.5% Triton X-100 in PBS for 10 min, and blocked in 2% BSA in PBS.

Antibodies to α-tubulin to stain microtubules or DRAQ5 to stain DNA were diluted in 1X PBS. After several washes with PBS, cells were incubated with the fluorescently tagged secondary antibodies for 1 h, washed in PBS and mounted with SlowFade®Gold antifade reagent (Thermo Fisher Scientific, Agawam, MA, USA).

### Microscopy

Microscopy studies were performed with a Nikon inverted TE2000-E microscope equipped with phase contrast and epifluorescence, a digital CoolSNAP HQ camera, a Prior ProScanII motorized stage and a Nikon C1 confocal system and with EZC1 and NIS-Elements acquisition software. Cytokinesis failure was quantified with 40X Nikon Cfi Plan Apo Dm oil objective.

### Cytokinesis assays

To assay the inhibition of cytokinesis, we quantified the number of cells that remained connected by midbodies (intercellular bridges) or that were binucleated after cells were stained with antibodies to α-tubulin and DRAQ5. Midbodies were identified by the high concentration of α-tubulin in both arms of the midbody (intercellular) bridge connecting presumptive daughter cells.

### Explant cultures

Mouse submandibular salivary glands were removed and microdissected from timed-pregnant female mice (Strain CD-1, Charles River Laboratories) at embryonic day E12-E13, with the day of plug discovery designated as E0. Experiments were performed in accordance with protocols approved by the IACUC committee at University at Albany, SUNY. Explants were cultured in single-well dishes (MatTek) on top of porous polycarbonate filters (Nuclepore, Whatman WHA110405) floating on 200 μm of serum-free DMEM/Ham’s F12 medium (F12) containing ascorbic acid (150 *μ*g/ml), transferrin (50 *μ*g/ml), 100 units/ml penicillin, and 100 μg/ml streptomycin, as reported previously (43–45).

### SiRNA transfection of salivary glands in *ex vivo* organ culture

Mouse submandibular salivary glands were dissected from timed-pregnant female mice (Strain CD-1, Charles River Laboratories) at embryonic day E12-E13. The day of plug discovery designated as E0. Experiments were performed in accordance with protocols approved by the University at Albany. Three rounds of overnight transfection using 400 nM siRNA with Oligofectamine were performed for optimal knockdown. After 88 h, ten non-targeting siRNA-treated glands and ten RSK2-siRNA treated glands were lysed and analyzed for RSK2 depletion and MKLP1 expression by western blot. An additional fifteen glands from each group were disrupted by dissociating the glandular buds with dispase and collagenase followed by gravity pellet enrichment for epithelial clusters as previously described (44,46). Dissociated cells were then plated onto a laminin-rich matrix (38,39) and immunostained with antibodies to α-tubulin and Hoechst or DRAQ5 to quantify binucleated cells.

### Statistical analysis

Statistical analysis was performed with GraphPad Prism software using a two-tailed Student’s t-test or one-way ANOVA with Dunnett’s post-hoc analysis. P value of p<0.05 was considered to be statistically significant.

## Acknowledgments

The authors thank Dmitri Wiederschain (Novartis Developmental and Molecular Pathways, Cambridge, MA) for generously providing the Tet-pLKO-puro lentiviral vector, and Debbie Moran for her help with the preparation of the manuscript.

